# Making honey bees lie: experimental dissociation of flight experience and dance communication

**DOI:** 10.1101/268292

**Authors:** Arumoy Chatterjee, M.V. Prabhudev, Ebi A. George, Pallab Basu, Axel Brockmann

## Abstract

Honey bees use their dance to communicate flight distance and direction of a food source to their nest mates in the hive. How bees transpose flight information to generate a corresponding walking (dance) behavior is still unknown. We now present a detailed study of the changes in dance duration of individual bees after shifting feeder distance. Our experiments indicated that most bees needed two or more foraging trips to the new position before showing an updated dance duration. In addition, only a few bees significantly changed dance duration immediately, whereas most bees first produced intermediary durations. Double shift experiments showed that under certain conditions bees do not update dance duration but continued to perform dance duration for the previously visited feeder position. We propose that generation of dance information involves two memory contents one for newly acquired and one for previously stored distance information.

**One Sentence Summary:** Generation of dance information is temporally separated from immediate flight experience and involves two different memory contents.

## Introduction

Honey bee foragers returning from a foraging trip communicate flight distance and direction to the food source to their nest mates using a small-scale walking pattern, the so-called waggle dance (*1*). Duration and orientation of the waggle run correlates with the flight distance and direction to the food source from the hive, respectively (*1–4*). Exploring how honey bees use flight information to produce waggle dances and vice versa is central for identifying neural mechanisms underlying dance communication. The time dynamics of these processes will provide insights about how tightly navigational and dance information processing are coupled.

In this study, we asked two questions: (i) How many foraging trips do honey bee foragers need to update their waggle dance duration? (ii) Do foragers use only information from the most recent foraging trip or do they also include information from earlier flight experiences to generate dance duration?

## Single shift experiment

First, we measured waggle dance durations for individually marked foragers visiting an unscented sugar-water feeder at a distance of 300m from the hive for 1-2 hrs. Then, the feeder was shifted for 100m, either forward (400m) or backward (200m) and we monitored the changes in dance behavior (Fig 1A; table S1; supplementary materials).

**Fig. 1.**
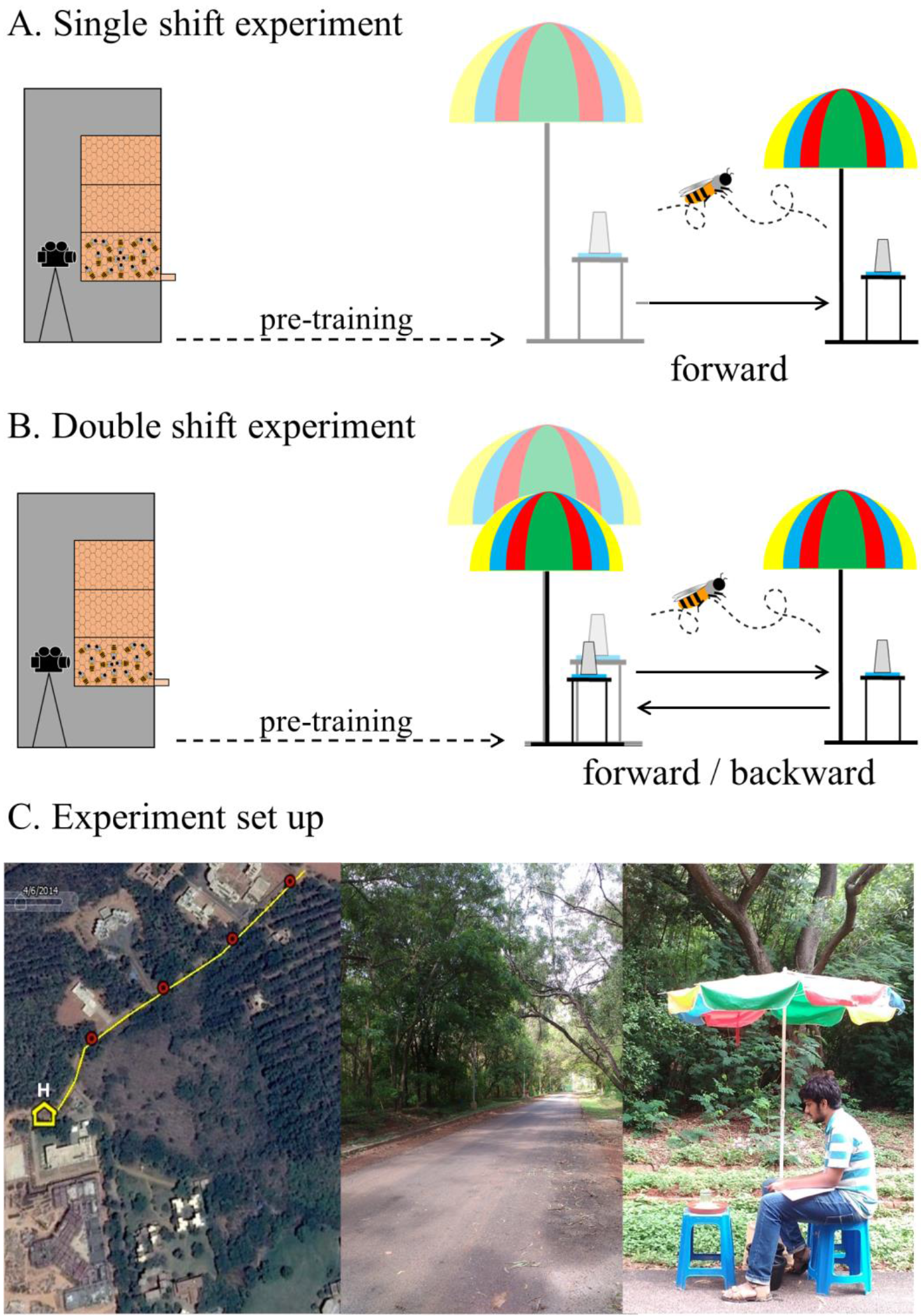
Honey bee foragers communicate the change in feeder location in waggle dance. (A) Single shift experiment: Foragers were trained to visit a feeder with non-scented sugar water 300m away from the hive (see also supplementary materials). During the experiment, individually marked foragers were allowed to visit the feeder for 1-2 hours; then the feeder was linearly displaced for 100m forward (400m) or backward (200m). (B) Double shift experiment: Similar experimental procedure as in the single shift experiment. In addition, the feeder was shifted back to the initial location after 1.5-2 hrs. (C) Google Image of the training path of the bees (left). H is the hive location and the red dots showed 100m, 200m, 300m and 400m feeder distances respectively. The training path offered necessary optic flow (middle). Feeder advertised for the bees with a multicolored umbrella (right).

Altogether, we trained 190 foragers (forward: n = 112, backward: n = 53), but only 35 individuals continued dancing after finding the feeder at the new distance. Fifteen out of these 35 foragers (43%;) continued dancing after the first visit, whereas 20 foragers (57%) stopped dancing for one or more foraging trips before they resumed dancing (Fig. 2A). We determined the first occurrence of a significant change in dance duration (= change point) for each individual (Fig. 2A-B; results of the change point analyses are provided in table S3). A total of 34 individuals showed a significant change-point in their post shift dance duration (table S3). Among the bees that continued dancing after the shift, 3 out of 15 (20%) foragers immediately showed a change point after the first feeder visit (= immediate update) while 12 foragers needed two or more foraging trips to update dance duration (= delayed update, Fig. 2B-C; table S3). Among the foragers that stopped dancing, 12 out of 19 (63%) foragers showed a change point in their first dance whereas 7 foragers still showed a delayed update (Fig. 2B; table S3). The proportion of bees showing change point in the very first post-shift dance was significantly higher in the group which stopped and resumed dancing (χ^2^-test: 63% vs 20%, χ^2^ = 3.84, df = 1, p = 0.03). The magnitude of change in dance duration was similar in the forward and backward shift (Fig. S2A).

**Fig. 2.**
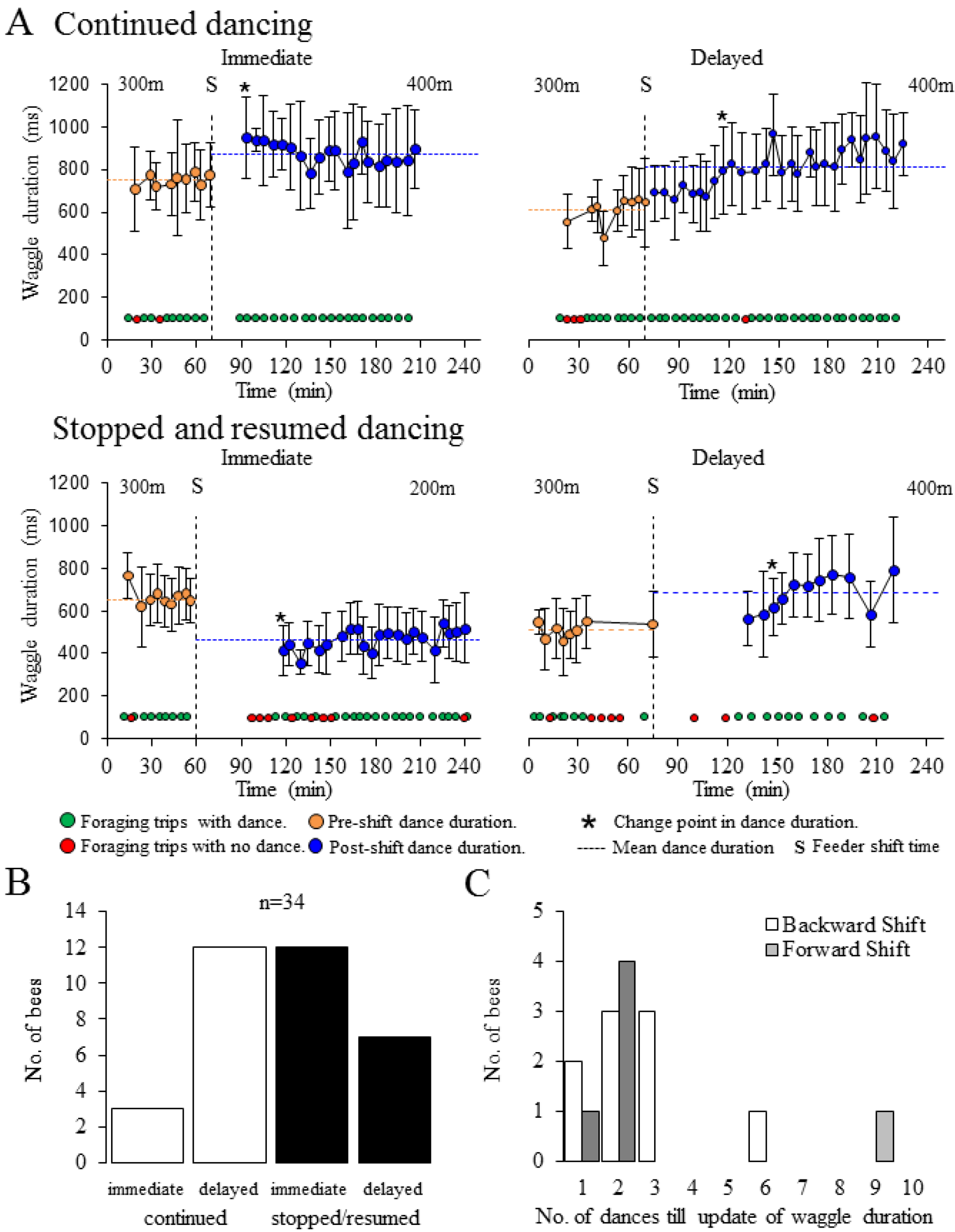
Single feeder shift experiments. A) After finding the new feeder location, foragers either continued dancing or first stopped and later resumed dancing. The change point determined the first post-shift dance with significantly different dance duration (supplementary materials). Individuals updated dance duration either on the very first post shift dance (immediate) or after one or more dances (delayed). Yellow and blue colored circles show pre-and post-shift dance duration. Error bars show standard deviation of waggle run duration. Green and red circles show feeder visit timings with or without following dances. Vertical dashed lines represent the timing of the feeder shift. Horizontal dashed line represents average dance duration before and after feeder shift (S). The asterisk indicates the change-point. B) 34 out of 35 foragers showed the change point on post-shift dance duration (table S3). We did not find any bias for post shift dance behavior (continued/stopped and resumed dancing, immediate/delayed update in dance duration) between forward or backward shift (χ^2^-test: χ^2^=0.682, df=3, p=0.8775). White bars for bees continuing dancing and black bars for those who stopped and resumed dancing, after feeder shift. C) 15 foragers continued dancing after finding the feeder at the new distance. Majority of foragers (12) needed multiple (>1) foraging trips to update dance duration, only 3 foragers updated dance-duration on the very first post shift dance. White and grey bars represents backward and forward shift respectively.

Further, we asked whether the change in dance duration occurred gradually or abruptly. A gradual update would include dances with intermediary durations (i.e. durations between the mean dance duration for pre-shift and post-shift dances). The occurrence of intermediary dances would indicate that bees use information not only from the most recent foraging trip but also from earlier foraging trips. We compared differences between consecutive dances in the pre-shift phase and post-change-point phase with those of the dances during the intermediary phase (= dance after the shift till the change point; supplementary materials).

Among the individuals which showed intermediary dances (n = 19), 15 (78.9%) of them showed a gradual change in dance duration, while 4 (21.1%) showed an abrupt change (Fig. S3A-C; analysis results in table S4). Within the group of bees that continued dancing, 10 out of 12 bees (83.3%) showed a gradual change and 2 out of 12 showed an abrupt change in dance duration (Fig. S3C; table S4). Second, we fitted two sigmoidal curves with different slopes to identify gradual and abrupt changes during the intermediary dances (supplementary materials, methods). Nine out of the total 19 (47.3%) bees showed a better fit with the gradual and 3 (15.7 %) a better fit with the abrupt model (Fig. S4). Six bees could not be grouped to either category. Among the group of continuously dancing bees, 6 out of 12 bees (50%) showed a better fit with the gradual model and 2 out of 12 showed a better fit with the abrupt model (Fig. S4; analysis results in table S4).

## Double Shift experiment

In a second set of experiments, we tested how honey bee foragers would adjust their dance behavior when confronted with shifting the feeder position twice, in opposite directions: either “backward-forward” (300m-200m-300m) or “forward-backward” (300m-400m-300m) (Fig. 1B; supplementary methods). Our question was whether or not the bees would show similar dance durations for the first and the second 300m feeder distance. Differences in the dance duration would indicate an effect of prior experiences.

In the backward-forward shift experiment, all foragers (n=11) showed significant changes in the waggle dance duration after the first and second shift (Fig. 3A-B). However, 7 out of the 11 foragers showed significantly shorter waggle run durations for the 300m(2) compared to the 300m(1) (number in brackets indicates first or second test at 300m distance; Fig. 3B, results from linear mixed effect modeling followed by general linear hypothesis testing are provided in table S6).

**Fig. 3.**
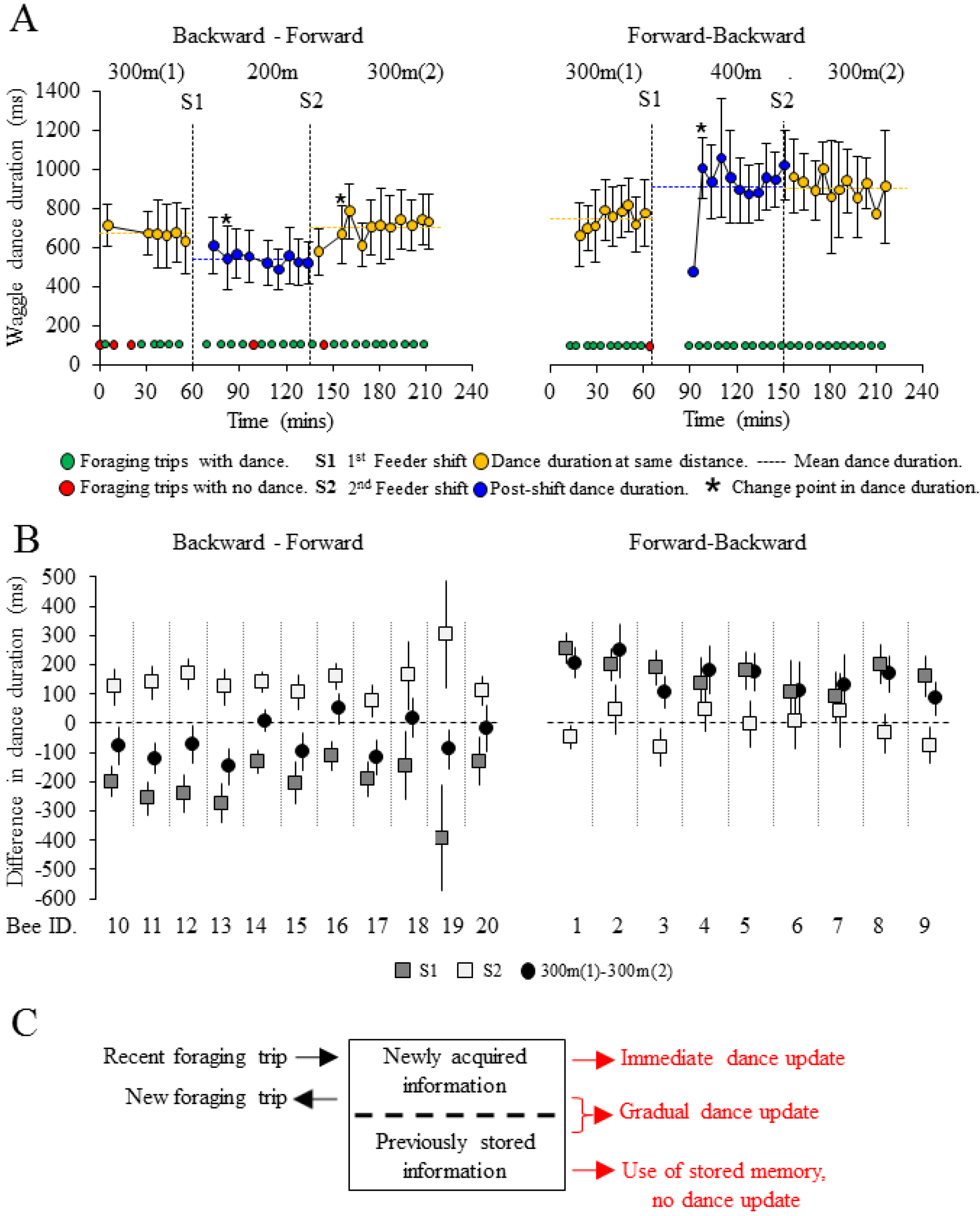
Double feeder shift experiments showed the role of past experience in shaping waggle dance duration. A) Individual forager updated dance duration (change point) following both feeder shifts (left) for backward-forward experiment but did not show any change in dance duration after the second shift (right) for forward-backward experiment (supplementary materials, table S5). Yellow colored circles show dance duration at same distance, blue circles show dance duration after first shift. Error bars show standard deviation of waggle run duration. Green circles show foraging trips followed by dances and red circles show same without dances. Horizontal dashed line represents average dance duration before and after first (S1) and second (S2) feeder shifts. The asterisk shows the change-points of dance duration. B) All foragers (n=11) showed significant change in dance duration after each shift (left) in the backward-forward experiments, yet 7 foragers (Bee ID. 10-13, 15,17,19) showed shorter dance duration while dancing for same distance (300m) after double feeder shift. While, in the forward-backward experiments, 8 foragers (n=9) updated dance duration after the first shift but none showed any change in dance duration after the second shift (right). Symbol shape represents difference in dance duration: solid rectangle = before and after first shift, hollow rectangle = before and after second shift and circles = same distance. Error bars show confidence intervals of the change in dance duration. C) Heuristic model of updating dance duration. We propose that flight navigational memory contains at least two different memory contents: (1.) *“newly acquired information”*, i.e. the most recent flight information, and (2.) *“previously stored information”*. In most cases both memories interact to guide dance behavior shortly after having found a new feeder position. During following trips, a new memory of *“previously stored information”* is generated. In the cases in which bees immediately updated they used the memory of *“newly acquired information”*. After the second feeder shift in the forward-backward experiment the bees communicated the *“previously stored information”* without updating dance information.

In the forward-backward shift experiment, 8 foragers (n = 9) showed a significant change in the dance duration after the first shift, but 6 foragers did not show any change in dance duration after the second shift (Fig. 3A-B). All foragers showed significantly longer dance duration for the first 300m(2) compared to the 300m(1). In fact, the dance durations for the 300m(2) were very similar to those of the previously visited 400m distance (−1.04 ± 2.82% change in dance duration, Fig. S5; table S7). We also did not find any change point after the second shift (Fig. 3A, table S5). An additional double shift experiment using different distances (forward-backward: 200m-300m-200m, n = 2) suggested that our finding is independent of the flight distances (Fig S6).

## Discussion

To summarize, our single shift experiments demonstrated that the majority of honey bee foragers needed multiple foraging trips to generate significantly changed, i.e. updated, dance duration. These results are supported by an observation by Tautz et al. (*5*), who mentioned, but did not analyze, a higher variability in waggle run duration during the first dances after a feeder shift. As there is ample evidence that ants and honey bees instantaneously know the walking or flight distance of a newly found food location (*6–8*), we propose that the additional trips (or additional time) are needed to exclusively generate the corresponding dance duration information. Thus, flight distance estimation and generation of waggle run duration are temporally separated.

Our double shift experiments provide two results. First, in the backward-forward experiment foragers showed shorter dance durations for the same feeder distance after feeder shifts. Second, in the forward-backward experiment bees did not change dance durations after the second shift. The forager basically ignored the new flight distance information and continued to perform the dance durations that they showed for the 400m. So far, the discussion on memory processes involved in generation of dance information have been focused on the sun-compass system and dance direction information (*9,10*). F. Dyer (*11*) proposed a model in which the generation of dance direction information involves two separate memories, a memory of a newly acquired navigational information from the most recent foraging trip and a memory of previously stored information from earlier flight experiences (Fig. 3C). The “newly acquired” memory will get incorporated in the “previously stored” memory, but both memories can be independently used to generate waggle dance duration. Some of the foragers in Dyer’s experiments showed an abrupt change in dance orientation ignoring previous experiences. Our forward-backward experiments showed foragers that did not change dance duration ignoring newly acquired navigational information. Similar to Dyer’s (*11*) ideas we propose that the generation of dance distance information is based on “newly acquired” and “previously stored” memory contents.

Recent radar tracking experiments showed that dance recruits compare newly received information from dancers with their own previous foraging experiences and use both information to decide where to search for food (*12*). We think that the results of our experiments point to a similar process in the dancers. First, the time delay between calculating flight distance and updating waggle run duration indicates that both information processing can be experimentally decoupled. Second, the double shift experiments showed that dancers are capable of generating dance information that is not related to the flight distance of the last foraging trip. Our results are supported by earlier studies that showed that foragers are able to perform appropriate dances during the night without a prior foraging trip (*13*). Together, all these experiments indicate that generation of dance information genuinely involves memory processes.

Dacke and Srinivasan (*14*) recently suggested that honey bee forager might have two different odometers, one for their personal use (i.e. flight navigation) and one for social (dance) communication. In our forward-backward experiments the foragers knew the correct flight distance otherwise they would not have continuously arrived at the 300m feeder position. However, for the communication of the distance they used their previously stored memory of the 400m feeder distance. We propose that both experiments report a similar phenomenon, if bees get confused or become uncertain about the distance of a feeder, they use the most recent navigational memory for their own orientation but communicate the previously stored (“confirmed”) navigational information to their nest mates.

Still the question remains why the foragers in the backward-forward experiment updated the waggle dance duration whereas the foragers in the forward-backward experiments did not. All earlier feeder shift experiments including the generation of distance calibration curves indicated that honey bees do not have any major difficulties with multiple forward or backward shifts of a feeder (*1,2,8,15*). If honey bees do not have any problem with double shift experiments, the difference in the behavior might have something to do with the spatial arrangement of the feeders or the order of feeder shifts. In the backward-forward experiments the new feeder position (200m) is actually on the way to the starting feeder position, whereas in the forward-backward experiments the new feeder position (400m) was not known before. We favor the idea that the novelty of the flight experience affected the response after the backward shift, but more experiments need to be done.

Finally, so far all attempts to identify molecular processes involved in dance communication failed because it was not possible to experimentally dissociate foraging flight and dance behavior (*16*). Our finding that most foragers need 2-3 foraging trips to update dance information opens a window to study molecular brain processes specifically involved in generating dance information (*17,18*).

## Authors’ Contributions

Experiments were designed by AC and AB and experiments were performed by AC and PMV. Dance analysis was done by AC. The statistical analyses were done by AC, EAG and PB. The paper was written by AB, AC and PB.

## Acknowledgments

We would like to thank B. Krishnan, S.K. Sethy, A. Sengupta, A. Suryanarayanan, S. Unnikrishnan, N. Thulasi, A. Johny, R. Fatima, A. Dey for their help with the behavioral experiments. We would like to thank UAS-GKVK, Bangalore for agreeing to let us use their campus. AC was supported by a fellowship from University Grants Commission; AB is supported by NCBS-TIFR institutional funds No. 12P4167.

